# A neonatal rat sepsis score captures the time course and severity of disease in a clinically relevant rat peritonitis model

**DOI:** 10.64898/2026.05.15.725467

**Authors:** Forough Jahandideh, Si Ning Liu, Kimberly B. Tworek, Ronan M.N. Noble, Jad-Julian Rachid, Aileen MacLellan, Manoj Lalu, Kimberly F. Macala, Stephane L. Bourque

**Affiliations:** Department of Anesthesiology & Pain Medicine, University of AB, Edmonton, Alberta, Canada; Women and Children’s Health Research Institute, University of AB, Edmonton, Alberta, Canada; Blueprint Translational Research Group, Acute Care Research Program, Ottawa Hospital Research Institute, Ottawa, ON, Canada; Department of Pediatrics, University of Alberta, Edmonton, AB, Canada; Department of Critical Care Medicine, University of Alberta, Edmonton, AB, Canada

**Author notes:** **Correspondence:** Stephane Bourque, PhD, Associate Professor, Departments of Anesthesiology & Pain Medicine and Pediatrics, Women and Children’s Health Research Institute, 3-020H Katz Group Centre; University of Alberta, Edmonton, Alberta, Canada T6G 2E1, E. These authors contributed equally: Forough Jahandideh and Si Ning Liu. **Category of study:** Basic science.

## Abstract

**Background:** Neonatal sepsis is a major cause of infant morbidity and mortality worldwide, particularly in preterm and very low birthweight babies. Fundamental differences between neonates and adults warrant clinically relevant models of neonatal sepsis. Here, we describe a preclinical fecal-slurry (FS)-induced peritonitis model of polymicrobial sepsis in neonatal rats, along with a novel neonatal rat sepsis score (nRSS) to monitor illness severity.

**Methods:** Peritonitis was induced in 3-day-old Sprague Dawley rats by intraperitoneal injection of various doses (0.3-1.5mg/g body weight) of fecal slurry (FS); control pups received equivalent doses of vehicle. All pups received analgesics (buprenorphine), antibiotics (ampicillin and gentamicin), and fluids (saline) to model clinical standards of sepsis treatment. Time-dependent changes in circulating cytokines (IL-6, IL-1β) and biomarkers of sepsis pathology (hemoglobin, glucose, alanine transaminase [ALT] levels) were assessed and correlated with nRSS scores.

**Results:** FS administration caused a dose-dependent increase in severity of sepsis over time, as indicated by increases in mortality rates (based on predefined criteria for euthanasia), nRSS scores, as well as time-dependent changes in circulating glucose, hemoglobin, IL-6, IL-1β, and ALT activity levels. nRSS scores correlated with all quantitative measures of sepsis pathology. Notably, females showed higher mortality and higher early NRSS scores than males at moderate to high FS doses, yet biochemical markers and time of death did not differ between sexes, suggesting that the apparent female vulnerability may reflect more conspicuous behavioral manifestations of illness rather than greater underlying physiological severity.

**Conclusion:** Induction of peritonitis in rats at postnatal day 3 produced a consistent and reproducible model of polymicrobial neonatal sepsis. Illness severity was monitored using a newly developed nRSS. By minimizing distress and incorporating standards of care, this model and scoring system may serve as a platform for future investigations into the underlying mechanisms and potential therapeutic interventions for neonatal sepsis.

**Impact:**

1. A clinically relevant rat model of neonatal polymicrobial sepsis was developed, incorporating standards of care (analgesics, antibiotics, and fluid resuscitation) to better reflect the clinical context in which preclinical findings must ultimately translate.
2. A novel neonatal rat sepsis scoring system (nRSS) was developed and validated, providing a sensitive, non-invasive measure of disease severity that correlates with biochemical markers and predicts mortality.
3. Female pups showed higher mortality and earlier behavioral signs of illness than males despite equivalent biochemistry, highlighting that clinical scores may capture sex-dependent vulnerability not apparent in standard biochemical measures.
4. Together, this model and scoring system offer a refined platform for mechanistic and therapeutic studies of neonatal sepsis while advancing the welfare-conscious 3Rs principles essential to rigorous preclinical research

## 1. Introduction

Neonatal sepsis is a life-threatening organ dysfunction resulting from a dysregulated host response to infection that occurs within 28 days of birth. It is commonly categorized as early-onset (occurring within 72h of birth, often caused by vertical transmission of pathogens from the mother during labor) or late-onset (occurring after 72h after birth, usually resulting from horizontal transmission from the surrounding environment). An estimated 1.3 million neonates are affected worldwide annually, with the highest incidence among premature and very low birthweight (<1,500 g) infants.^1^ The global incidence is expected to rise with increasing rates of premature births.^2^ Mortality rates depend on patient vulnerability and causative pathogen, but case fatality ratios of 10–35% are typically reported.^3-5^ Among surviving infants, neonatal sepsis remains a major determinant of poor neurodevelopmental outcomes and chronic health complications.^6,7^

The neonate exhibits distinct innate and adaptive immune responses and has unique metabolic demands that differ from those of older children and adults.^8,9^ These differences likely underlie divergent clinical trajectories observed between neonatal and adult sepsis, including distinct patterns of organ dysfunction, differential susceptibility to specific pathogens, and differing responses to therapeutic interventions.^8,9^ Despite this, neonatal sepsis has historically been understudied relative to adult sepsis, and much of our mechanistic understanding has been extrapolated from studies in adults.^10^ This extrapolation is a critical gap: interventions that show promise in adult sepsis models frequently fail in the neonatal context,^11-13^ highlighting the need for neonatal-specific research frameworks.

Effective therapies that directly address the underlying pathophysiology of neonatal sepsis-induced organ dysfunction are lacking. Current management strategies primarily involve pathogen clearance (e.g., source control and antibiotic therapy) and supportive measures. The dearth of promising new interventions is partly attributed to an inability to validate findings from preclinical animal models in clinical studies.^14^ Preclinical study designs that fail to recapitulate the clinical condition of sepsis likely contribute to the poor translatability.^14^ The Minimum Quality Threshold in Pre-Clinical Sepsis Studies (MQTiPSS) recommendations, developed through the Wiggers-Bernard Conference and representing consensus from 31 experts from 13 countries, provides guidelines to standardize preclinical sepsis models and improve translational relevance.^15^ These guidelines address key experimental design elements, including model relevance to clinical conditions, monitoring criteria for distress and euthanasia, and use of analgesics and antimicrobial agents.^14,16^ However, existing MQTiPSS-aligned models have been developed primarily in the context of adult sepsis, leaving a significant gap in standardized, clinically relevant preclinical tools for neonatal diseases.

Rodents are the most widely used mammalian models for sepsis research. Methods to induce sepsis in rodents include single-pathogen administration into the blood, brain, lung, gastrointestinal system, or peritoneal cavity, as well as polymicrobial intraperitoneal challenge.^17^ The fecal slurry induced peritonitis (FIP) model of polymicrobial sepsis was initially described in adult rats,^18^ and has been applied in neonatal mice^19-22^ to model clinical conditions such as necrotizing enterocolitis or spontaneous bowel perforation, which occur primarily in premature and extremely preterm infants.^23,24^ However, neonatal mice present practical limitation that constrain the depth of mechanistic investigation: their small body size restricts tissue availability for biochemical analyses, and dams are prone to cannibalizing pups during stress – a significant concern for animal welfare and experimental reproducibility.

To address these limitations and advance neonatal-specific research, we developed and characterized a neonatal rat FIP model that incorporates standards of clinical care, including analgesics, antibiotics, and fluid resuscitation in alignment with MQTiPSS guidelines. The larger body size of neonatal rats provides distinct advantages over mice, including greater tissue yields for biochemical analyses, and improved feasibility for complex invasive procedures. Moreover, larger litter sizes enhance experimental throughput and statistical power. We utilized postnatal day-3 rats, a developmental stage that closely approximates human prematurity.^25,26^ To enable objective, real-time monitoring of disease progression, we also developed and validated a novel neonatal Rat Sepsis Score (nRSS) for assessing illness severity. The nRSS was specifically designed to minimize the duration of maternal separation – a critical welfare consideration in neonatal rodent studies – while providing reliable and quantifiable assessment of sepsis severity.

## 2. Materials and Methods

All experimental protocols were approved by the University of Alberta Animal Care and Use Committee in accordance with the guidelines of the Canadian Council of Animal Care. This study was conducted and reported in accordance with the Anima Research: Reporting of In Vivo Experiments (ARRIVE 2.0) guidelines.^27^

### 2.1. Fecal slurry preparation

Fecal slurry (FS) was prepared as previously described^28^ with minor modifications. Briefly, cecal contents were harvested from untreated, humanely euthanized male Sprague Dawley rats (10-12 weeks of age) and homogenized in 50 mM potassium phosphate buffer. The homogenate was filtered through a 100 µm cell strainer to remove large particles, and then centrifuged at 3000 × g for 25 min at 4°C. The resulting pellet was resuspended in 5% dextrose and 10% glycerol in ddH_2_O to a final concentration of 100 mg/mL. Aliquots were flash frozen in liquid nitrogen and stored at -80 °C; each aliquot was used after a single freeze-thaw cycle.

### 2.2. Animals and treatments

Male and female Sprague Dawley rats (10-12 weeks of age) were purchased from Charles River (K98; Kingston, NY) and housed in same-sex pairs at the University of Alberta animal care facility under a 12-h light/dark cycle [07:00-19:00] at 23°C and 40–60% relative humidity, with *ad libitum* access to a standard rodent chow (5L0D; PicoLab, St Louis, MO, USA) and water. After 1-2 weeks of acclimation, females were bred by overnight by co-housing with age-matched males. Pregnancy was confirmed by the presence of sperm in vaginal smears the following morning, after which females were single housed for the duration of the study. Within 12h of giving birth (defined as postnatal day (PD)0), litters were culled to 12 pups to standardize postnatal conditions.

At PD3, rat pups were allocated to control or one of five FIP dose groups by sequential alternating assignment within each litter, ensuring balanced litter representation across groups. FIP pups received an intraperitoneal injection of FS at T=0h (T0h) at one of five doses (0.3, 0.6, 0.9, 1.2, and 1.5 mg/g·body weight (BW); control pups received an equivalent volume of vehicle (Veh; 5% dextrose and 10% glycerol in ddH_2_O). All pups received a subcutaneous injection of slow-release buprenorphine (0.5 mg/kg·BW; Chiron Compounding Pharmacy Inc. Guelph ON) at T0h for analgesia; the slow-release formulation was selected to minimize the number of injections, handling, and disruption of dam-pup interactions. Antibiotic therapy consisted of subcutaneous ampicillin (20mg/kg BW) and gentamicin (4mg/kg BW), both administered at T4h, with an additional dose of ampicillin at T16h; antibiotic selection was guided by FS culture and sensitivity results (Prairie Diagnostic Services Inc., Saskatoon, SK). Fluid resuscitation (sterile saline) was administered subcutaneously at T4h and T16h, with total daily fluid volume standardized to 5mL/kg BW/day across all pups. Pup health was monitored every 2h starting at T4 until T16, and then every 4h until T24h (**Fig. 1**).

**Figure 1.**
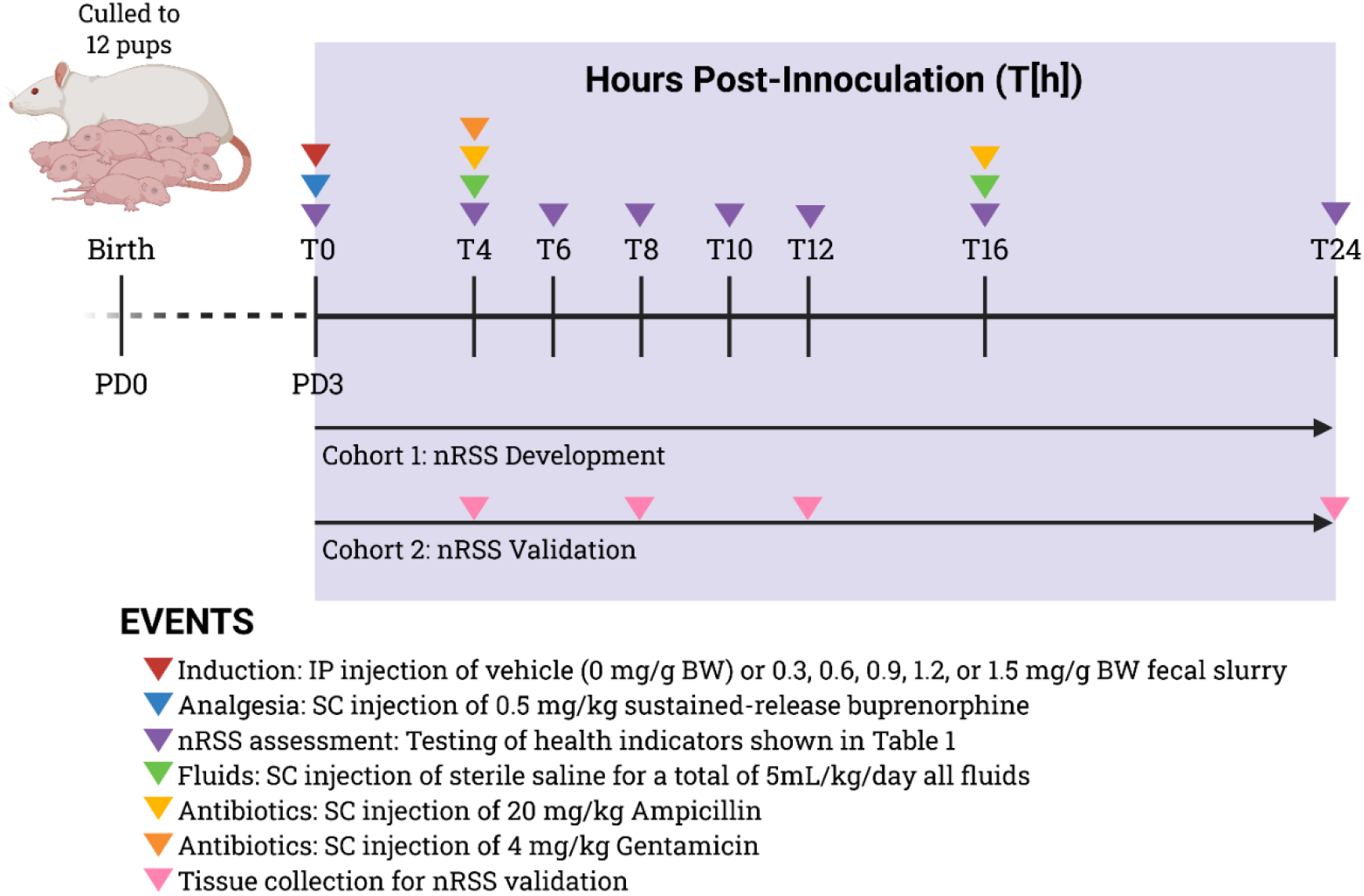
Experimental timelines for cohorts 1 and 2. Litters were culled to 12 pups per litter within 12 hours of birth. Coloured wedges denote interventions after fecal-slurry or vehicle (5% dextrose in 10% glycerol solution) administration (defined as time=0h, or T0h). BW: body weight, IP: intraperitoneal, nRSS: neonatal rat sepsis score, PD: postnatal day, SC: subcutaneous. Created with Biorender.com.

### 2.3. Development of neonatal rat sepsis scoring system

Candidate health indicators for the nRSS were initially drawn from published scoring systems for adult^29^ and neonatal septic mice,^30,31^ and included parameters related to appearance (skin color, presence of milk spot), pain or distress (hunched posture), spontaneous movement (mobility, righting reflex) and provoked behavior (response to tactile stimulation). Additional parameters included effort to rejoin the litter after separation, breathing pattern, and skin temperature measured by infrared thermometry (Flash-III; TLD Inc., Dawsonville, GA, USA) (**Suppl. Table S1**).

In keeping with refinement principles for animal-based research, humane endpoints were used as a proxy for mortality to minimize suffering. Humane endpoint criteria were defined in consultation with a clinical veterinarian (DD) based on respiratory distress or abnormal breathing pattern (laboured breathing, cyanosis, markedly altered respiratory rate), severe lethargy (minimal movement or inability to ambulate), or loss of limb movement. Pups meeting any of these criteria were promptly euthanized and classified as non-survivors. All mortality reported herein therefore reflects humane euthanasia rather than spontaneous death.

**Suppl. Table S1.** Parameters considered for developing the neonatal rat sepsis score (nRSS).

| Parameter | Levels | Description |
| --- | --- | --- |
| Regrouping | Yes/No | Pup rejoins litter after separation from dam and other pups |
| Skin temperature (°C) | Numerical | Taken within 30s of removing dam from the cage |
| Skin color | Pink/Pale/Grey | Judged in comparison to other pups within the litter |
| Breathing pattern | Irregular/Labored | Increased or decreased respiratory rate, respiratory distress, labored breathing. |
| Milk spot | Present/Absent | Presence of milk in the stomach |
| Hunched posture | Present/Absent | Presence of obvious hunch, which tends to be accompanied by the tail curled underneath the body. |
| Mobility | Active/Inactive | Any movement away from the pup's original location |
| Righting reflex | Able to right/Fail to right | Ability to right when pups are placed supine. Given 2 attempts (60s each). Also consider the effort expended if pup fails to right. |

In the first cohort (n=117 pups from 10 litters), all nRSS parameters were assessed at each monitoring time point (**Fig. 1)** until a humane endpoint was reached or until T24h, at which point all surviving pups were euthanized. In the second cohort (n=305 pups from 25 litters), pups underwent identical treatments but were euthanized by decapitation at predetermined experimental endpoints of T4h, T8h, T12h or T24h for tissue collection. Pups reaching humane endpoints prior to their assigned experimental endpoint were euthanized and classified as non-survivors. Tissues were not collected from pups reaching humane endpoints.

### 2.4. Blood glucose, hemoglobin, and plasma analysis

Following pup decapitation, trunk blood was assessed for glucose using a Contour Next EZ glucometer and for hemoglobin using a HemoCue Hb201+ hemoglobinometer. Remaining blood was collected into EDTA-coated tubes (Microvette CB300 K2E, Sarstedt AG & Co. KG, Numbrecht, Germany) and centrifuged at 1,500 × g for 15 min at 4°C. Isolated plasma was flash frozen in liquid nitrogen and stored at −80°C until analysis. Plasma concentrations of interleukin (IL)-1β and IL-6, markers of systemic inflammation, were quantified by enzyme-linked immunosorbent assays (ELISA) using commercially available kits (RLB00 and R6000B, respectively; R&D systems, Minneapolis, MN, USA) per the manufacturer’s instructions. Plasma alanine aminotransferase (ALT) activity, a marker of hepatic injury, was measured using a colorimetric assay kit (MAK052, Sigma Aldrich, St. Louis, MO, USA) per the manufacturer’s instructions. Each pup contributed a single value at its assigned endpoint (4, 8, 12, or 24 h).

### 2.5. Statistical analyses

Time-to-event analyses used Cox proportional hazards regression with a random effect for dam to account for litter clustering (dam-level mortality SD = 1.32 on the log-hazard scale). Pups euthanized at defined tissue-collection endpoints (T4h, T8h, T12h, T16h, or T24h) were treated as right-censored; pups that died spontaneously or were euthanized after reaching humane endpoints (nRSS ≥ 9) were treated as events at their time of death. FS doses were grouped a priori as low (0– 0.9 mg/g), mid (1.2 mg/g), and high (1.5 mg/g). Dose × sex interaction was assessed by likelihood ratio test. Marginal Cox models for sex and pup weight were compared by AIC to identify the dominant predictor. Results are reported as hazard ratios (HR) with 95% CIs.

Dose effects on total nRSS score, nRSS components (righting, colour, mobility), and biochemical markers were assessed using linear mixed models, with sex and time as fixed effects and dam as a random effect to account for litter clustering. Intraclass correlation analysis confirmed substantial dam-level clustering for nRSS (median ICC ≈ 0.09) and biochemical markers (median ICCs 0.10–0.19). All-pairwise dose comparisons were adjusted by Tukey’s HSD. The prognostic value of nRSS for eventual non-survival was assessed in two subsets: all FS-treated pups (0.3–1.5 mg/g) and the mid FS dose group (1.2 mg/g); vehicle-treated pups were excluded. Receiver operating characteristic (ROC) curves were generated at T4h, T6h, and T8h. Primary analysis of plasma biochemical markers included only survivors; non-survivor values are presented descriptively. ALT was log-transformed and IL-6 and IL-1β were log(x+1)-transformed prior to analysis; all other data were analyzed as raw values.

The relationship between nRSS scores and biochemical markers was assessed using Spearman rank correlations among surviving pups, including vehicle controls (n=131-268 per marker). Each surviving pup contributed biomarker values only at the time of euthanasia, paired with nRSS scores recorded at the same timepoint (concurrent nRSS) for markers whose disturbances resolved by T24h (glucose, Hb, IL-6), or with the highest nRSS recorded (maximum nRSS) for markers whose disturbances persisted beyond T24h (ALT, IL-1β). ALT, IL-6, and IL-1β were log(x+1)-transformed for plotting and bin-mean computation due to right-skewed distributions and zero values. Glucose was expressed as the absolute deviation from the vehicle-control median (6.4 mmol/L) to capture dysregulation magnitude regardless of direction, given the biphasic time course of glucose changes during sepsis. Non-survivors are overlaid at their maximum nRSS bin as separate symbols and compared to the highest-severity survivor pool (nRSS ≥ 6) by Mann-Whitney U test. Finally, as a sensitivity analysis, per-timepoint Spearman correlations between concurrent nRSS and each biochemical marker were computed at 4, 8, 12, and 24 h.

Finally, interobserver reliability for nRSS scoring was assessed using Gwet’s AC2 coefficient with quadratic weights. Statistical analyses were performed using Prism 10 (GraphPad Software Inc., San Diego, U.S.A) or in R (version 4.5.3).

## 3. Results

### 3.1. Dose-dependent mortality in the neonatal FIP model

Of 493 pups, 87 (17.6%) died or required humane euthanasia during their observation window (administrative censoring at the assigned endpoint, T4h-T24h). Mortality rose steeply with dose: Kaplan-Meier-estimated mortality 24h mortality was 2.2% in the low FS group (0–0.9 mg/g, n=265), 43.0% in the mid group (1.2 mg/g, n=182), and 67.4% in the high group (1.5 mg/g, n=46) (**Fig. 2A**; Cox dose effect LR χ^2^ = 84, P<0.0001). Litter clustering was substantial (random effect SD=1.32), indicating shared risk among pups from the same dam independent of dose.

**Figure 2.**
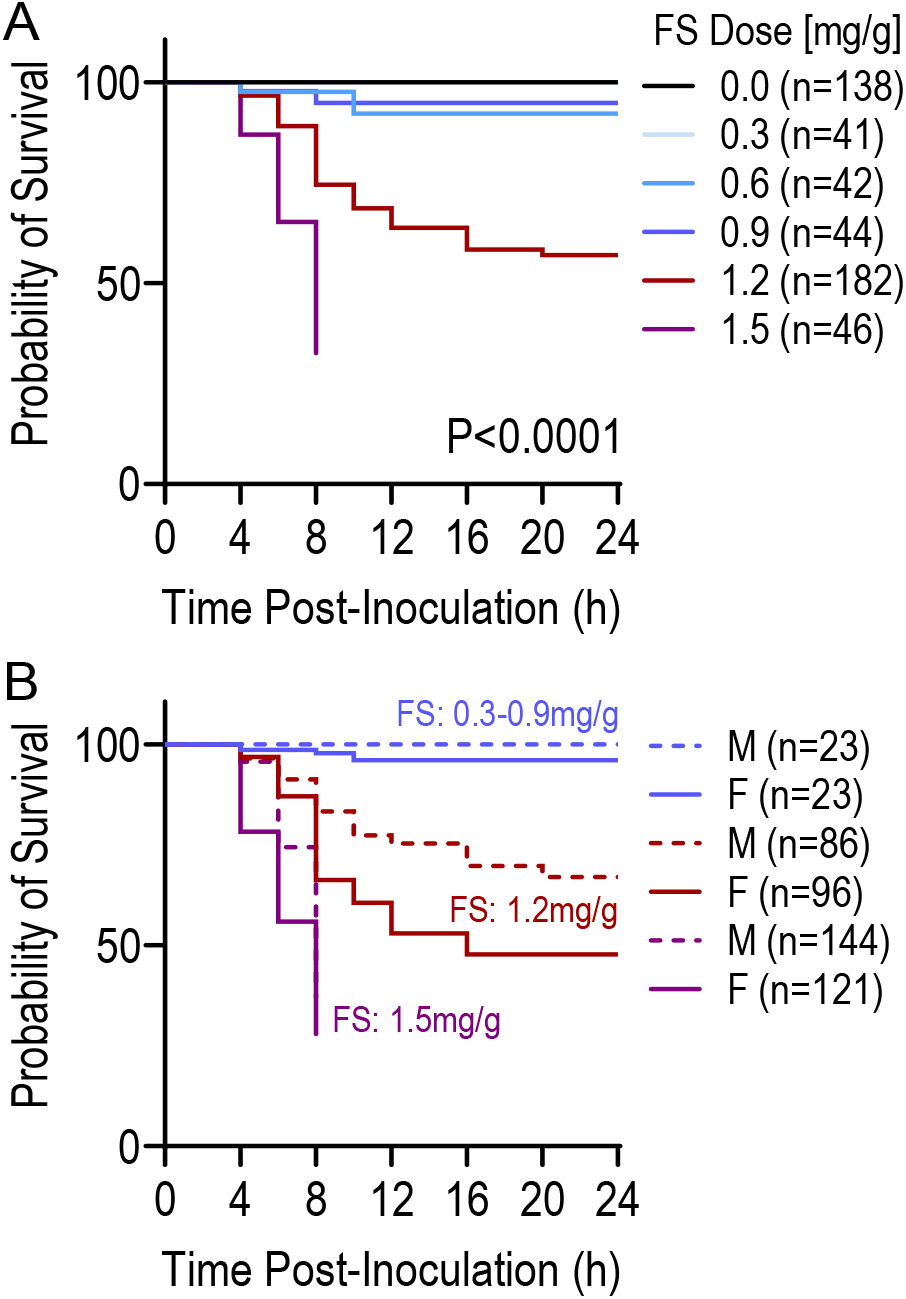
Mortality in FS-induced peritonitis is dose dependent and modified by sex in neonatal rats. (A) Kaplan-Meier survival curves for all pups (males and females combined) following FS inoculation on postnatal day 3, stratified by FS doses: Low (0.3-0.9mg/g; blue), Mid (1.2mg/g; red) and High (1.5mg/g; purple) (B) Survival curves stratified by both dose group FS dose groups and sex (solid lines, females; dashed lines, males). Time-to-event analyses were performed by Cox proportional hazards regression with a random effect for dam to account for litter clustering; see Methods for full statistical details.

A significant dose × sex interaction emerged in the full Cox model (LR χ^2^=8.05, P=0.018). In the mid-dose group, females had higher mortality than males (38.5% vs 23.3% mortality; HR=0.53, 95% CI 0.29–0.97, P=0.038; **Fig. 2B**), and a similar pattern was evident in the high dose group (60.9% vs 52.2%; HR=0.35, 95% CI 0.14–0.88, P=0.026). To examine whether pup body weight also modified mortality risk independent of dose, 1.2mg/g FS dose pups were stratified into weight tertiles (low: <9.24g; Mid: 9.25-10.00g; High>10.01). Mortality differed substantially across tertiles (Cox LR χ^2^=6.55, P=0.038): pups in the highest tertile had markedly lower mortality (14.8%) than those in the lowest (41.0%) or middle (38.3%) tertiles (**Suppl. Fig. 1A**). However, because birth weight and sex were collinear (males were heavier than females: 9.87g vs 9.40g, P<0.001) and thus males constituted a higher proportion of the highest body weight tertile (**Suppl. Fig. S1B, C**) univariate Cox models were compared to identify the dominant predictor. The sex-only model fit better than the weight-only model (ΔAIC=-3.3). In a combined model, the effect of sex remained after adjustment for weight (HR 0.53 → 0.54), whereas the effect of weight was abolished after adjustment for sex (HR per gram 0.78 → 0.97; P=0.90). Thus, the apparent protective effect of higher pup weight reflects confounding by sex. Among pups that died, time of death did not differ between sexes (p = 0.19).

**Supplementary Figure S1.**
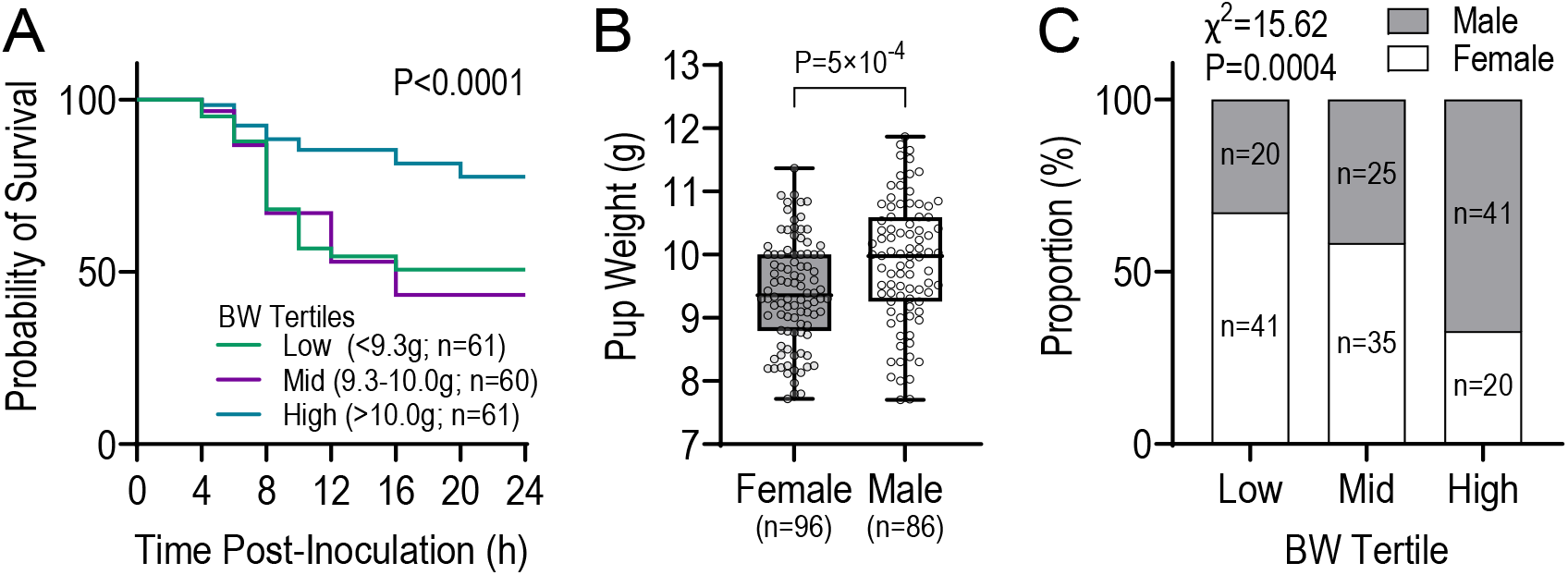
The apparent effect of body weight on mortality reflects confounding by sex. (A) Kaplan-Meier survival curves for pups in the Mid FS group (1.2mg/g) stratified by body weight (BW) tertile (Low, Mid, High). (B) BW distribution by sex within the Mid FS group, showing males are heavier than females (Welch’s *t*-test, P=0.0005). (C) Sex distribution within each BW tertile, showing systematic enrichment of females in the lower tertile and males in the higher tertile (χ^2^ test, P=0.004).

### 3.2. Development of the nRSS

To develop the nRSS, we systematically evaluated phenotypic and behavioural parameters and assigned scoring ranges that captured biological variance on a scale amenable to rapid assessment. Pilot studies identified skin color, righting ability/effort, and mobility as the parameters most consistently correlated with survival (**Table 1)**. We then assessed individual parameters and their combinations – righting + color, righting + mobility, color + mobility, and righting + color + mobility) – based on their ability to discriminate between pups that did and did not meet humane endpoint criteria (**Table 2**). Mobility and color alone had limited discriminatory power. Righting score was the strongest single predictor but was associated with a 29% overlap between survivors and non-survivors – meaning that use of righting score alone would result in unnecessary euthanasia for nearly one-third of non-moribund pups. The addition of color and/or mobility scores progressively improved discrimination, and the sum of all three parameters yielded the lowest overlap (8.5%) (**Suppl. Table S2**). Sepsis severity categories were therefore defined based on the cumulative nRSS (**Table 1**): pups scoring above 7 were at high risk of progressing to severe sepsis requiring euthanasia, while scores of 9 or 10 at any point after T4h indicated a moribund state requiring immediate euthanasia. Based on these thresholds, the nRSS demonstrated 89.4% between agreement with clinical judgement (i.e. reaching humane endpoints) to identify pups requiring euthanasia.

**Table 1.**
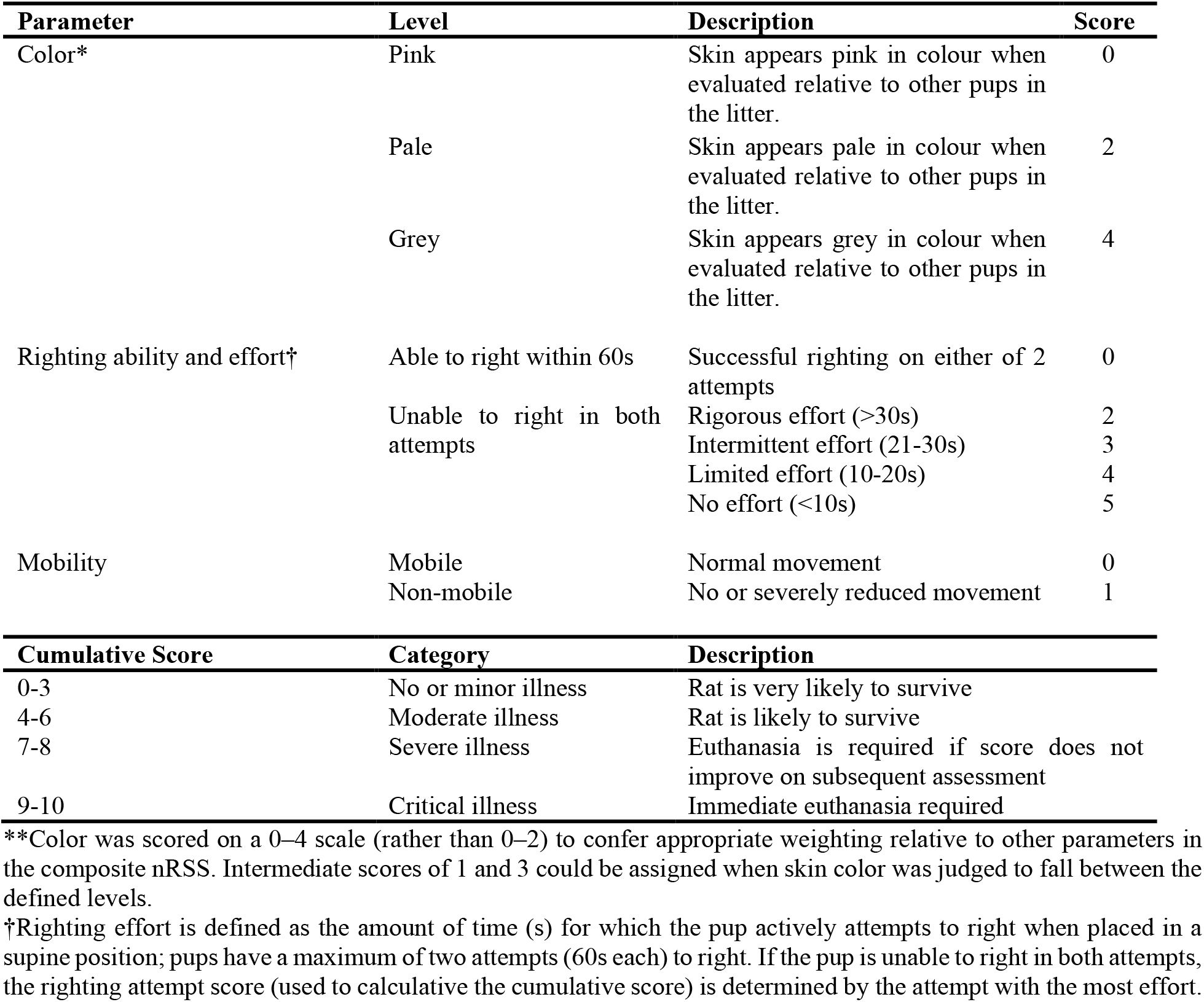
Neonatal rat sepsis score (nRSS) parameters and corresponding scores.

Two trials were conducted to assess interobserver reliability of the nRSS in pups administered either vehicle or FS; the first trial consisted of three blinded observers (one experienced and two naïve to the scoring system), and the second trial consisted of 4 blinded observers (two experienced and two naïve). Unexperienced observers received written criteria and completed a single training session the day prior to assessment. Pups were evaluated concurrently by all observers at T4h, T6h and T8h (n=12 pups in trial 1; n=24 pups in trial 2), without conferral between observers, to minimize repeated handling and litter disruption. Agreement among observers was high in both trials (trial 1: AC2=0.91, 95% CI: 0.88–0.95; trial 2: AC2=0.93, 95% CI: 0.91-0.96).

**Suppl. Table S2.** Specificity of neonatal rat sepsis score (nRSS) parameters and their combination on distinguishing between surviving and non-surviving pups.

| Parameters | Groups | Score range | Overlap range | Overlap (n/N) | Overlap % |
| --- | --- | --- | --- | --- | --- |
| Righting | Survivors | 0-5 | 2-5 | 34/117 | 29.1% |
|  | Non-survivors | 2-5 |  |  |  |
| Color | Survivors | 0-4 | 2-4 | 83/117 | 70.9% |
|  | Non-survivors | 2-4 |  |  |  |
| Mobility | Survivors | 0-1 | 0-1 | 117/117 | 100% |
|  | Non-survivors | 0-1 |  |  |  |
| Righting+Color | Survivors | 0-9 | 6-9 | 15/117 | 12.8% |
|  | Non-survivors | 6-9 |  |  |  |
| Righting+Mobility | Survivors | 0-6 | 3-6 | 18/117 | 15.4% |
|  | Non-survivors | 3-6 |  |  |  |
| Color+Mobility | Survivors | 0-5 | 2-5 | 84/117 | 71.8% |
|  | Non-survivors | 2-5 |  |  |  |
| Righting+Color+Mobility | Survivors | 0-9 | 7-9 | **9/117** | **8.5%** |
|  | Non-survivors | 7-10 |  |  |  |

### 3.3. nRSS captures dose dependent illness and predicts eventual mortality

FS administration produced dose-dependent rise in nRSS at all timepoints assessed (overall dose effect P<0.0001 from T4h–T16h; P=0.011 at T24h; **Fig. 3A**). Responses showed three distinct groupings: (i) controls (0 mg/g), which remained unchanged throughout; (ii) low dose FS group (0.3–0.9 mg/g) formed a low-elevation cluster with no pairwise differences at any timepoint; and (iii) mid-high FS dose (1.2–1.5 mg/g) groups formed a higher-elevation cluster that differed only at T8h (P=0.005). At peak illness (T8h), 12 of the 15 Tukey-adjusted pairwise comparisons in nRSS between doses were significant, with only non-significant comparisons being those among the three lowest doses (0.3, 0.6 and 0.9mg/g BW). The same pattern was preserved across all three nRSS components (righting, colour, and mobility; **Fig. 3B-D**). In the mid dose group (1.2mg/g), females had higher nRSS than males during the early phase (T4–8h, all P<0.025), indicating that the vulnerability in females is detectable by nRSS before mortality occurs.

**Figure 3.**
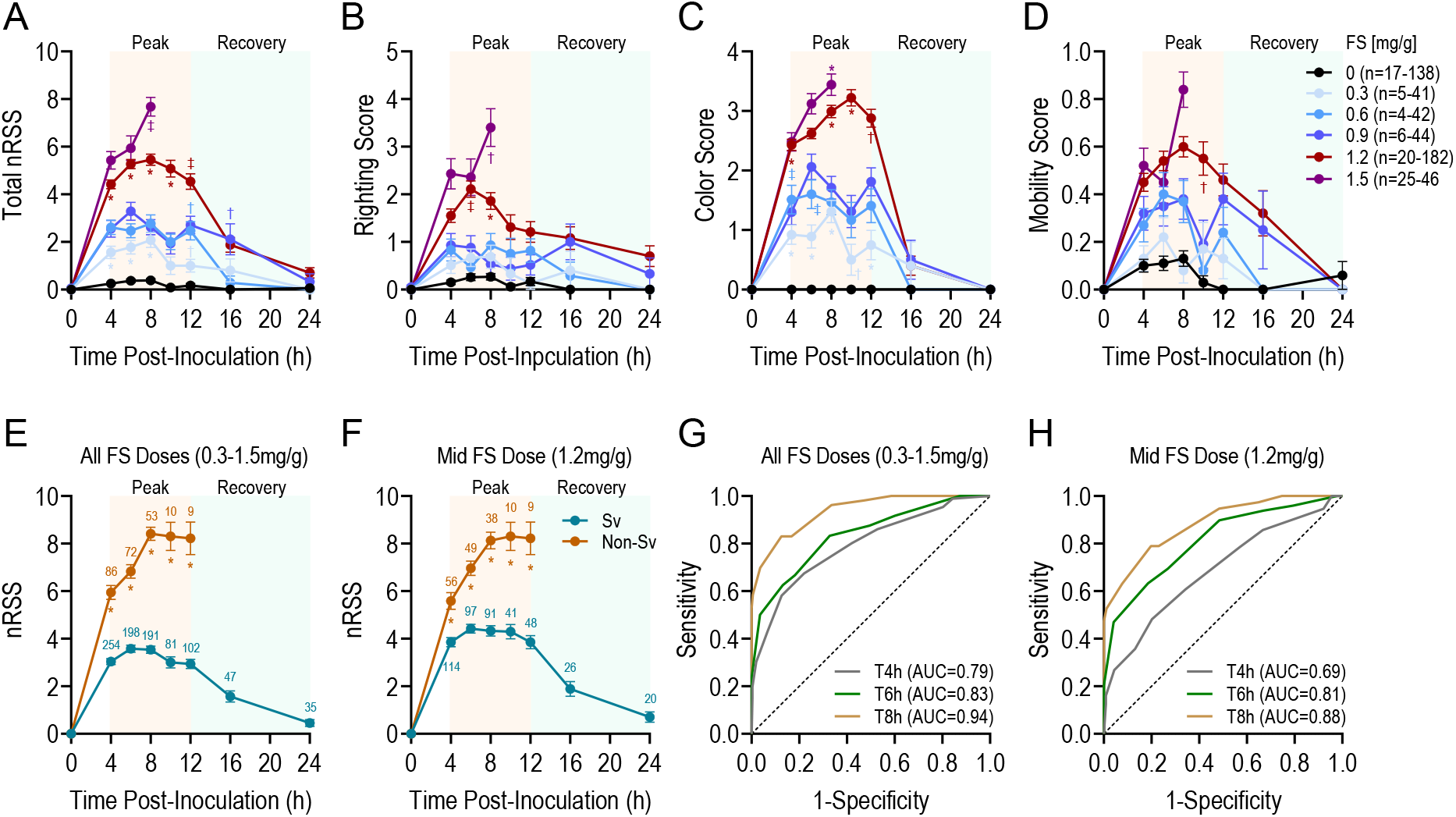
nRSS captures the time course of sepsis severity and predicts mortality. (A) Dose- and time-dependent changes in total nRSS following FS-inoculation, showing distinct peak and recovery phases across doses. (B-D) Individual component scores: (B) righting, (C) color, and (D) regrouping behaviour. (E, F) nRSS profiles in surviving (Sv) and non-surviving (non-Sv) pups across all challenged groups (0.3-1.5mg/g; E) panel) and within the Mid FS group along (1.2mg/g; F), illustrating the divergent trajectories that distinguish pups destined to survive from those approaching humane endpoints. (G, H) Receiver operating characteristic (ROC) analyses showing the predictive ability of nRSS scores for mortality at successive timepoints, in all challenged groups combined (0.3-1.5mg/g; G) and within the mid FS group alone (1.2mg/g; H). Data in A-F are mean±SEM; numbers above points in E-F denote n values.

When pups were stratified by eventual survival outcome, nRSS trajectories diverged sharply within hours of FS administration. Among all FS-challenged pups (0.3–1.5 mg/g), non-survivors had higher nRSS than survivors as early as T4h, and gap widened through T8h (**Fig. 3E**); the same pattern was evident in the mid FS dose group (**Fig. 3F**). By T8h, mean nRSS in non-survivors approached the humane endpoint threshold (nRSS≥9), while survivors plateaued at substantially lower scores. nRSS measured at early post-inoculation timepoints was strongly predictive of eventual non-survival. Among all FS-challenged pups, the AUC increased from 0.79 at T4h to 0.94 by T8h (**Fig. 3G**). nRSS retained substantial discriminative ability in the mid FS dose alone (AUC=0.69, 0.81, and 0.88 at T4h, T6h, and T8h, respectively; **Fig. 3H**).

### 3.4. Sepsis disrupts metabolic and immune function in neonatal rats

In response to FS challenge, glucose showed a striking biphasic response: at T4h, FS-challenged pups exhibited dose-dependent hyperglycemia, with mean glucose reaching 24.9 and 26.1 mmol/L in the 1.2 and 1.5 mg/g groups respectively, compared with 6.5 mmol/L in vehicle-treated pups (overall dose effect P<0.0001; **Fig. 4A**). By T12h, this pattern reversed: all surviving FS-challenged pups were hypoglycemic (mean glucose 2.2–4.2 mmol/L vs 6.5 in controls; P<0.0001), reflecting the transition from acute stress hyperglycemia to fuel exhaustion. Hb levels were elevated in a dose-dependent manner in FS-challenged pups during the early-to-mid phase (Fig 4B; P<0.001 at T4h, T8h, and T12h), consistent with hemoconcentration from fluid shifts; this resolved by T24h (P=0.13; **Fig. 4B**).

**Figure 4.**
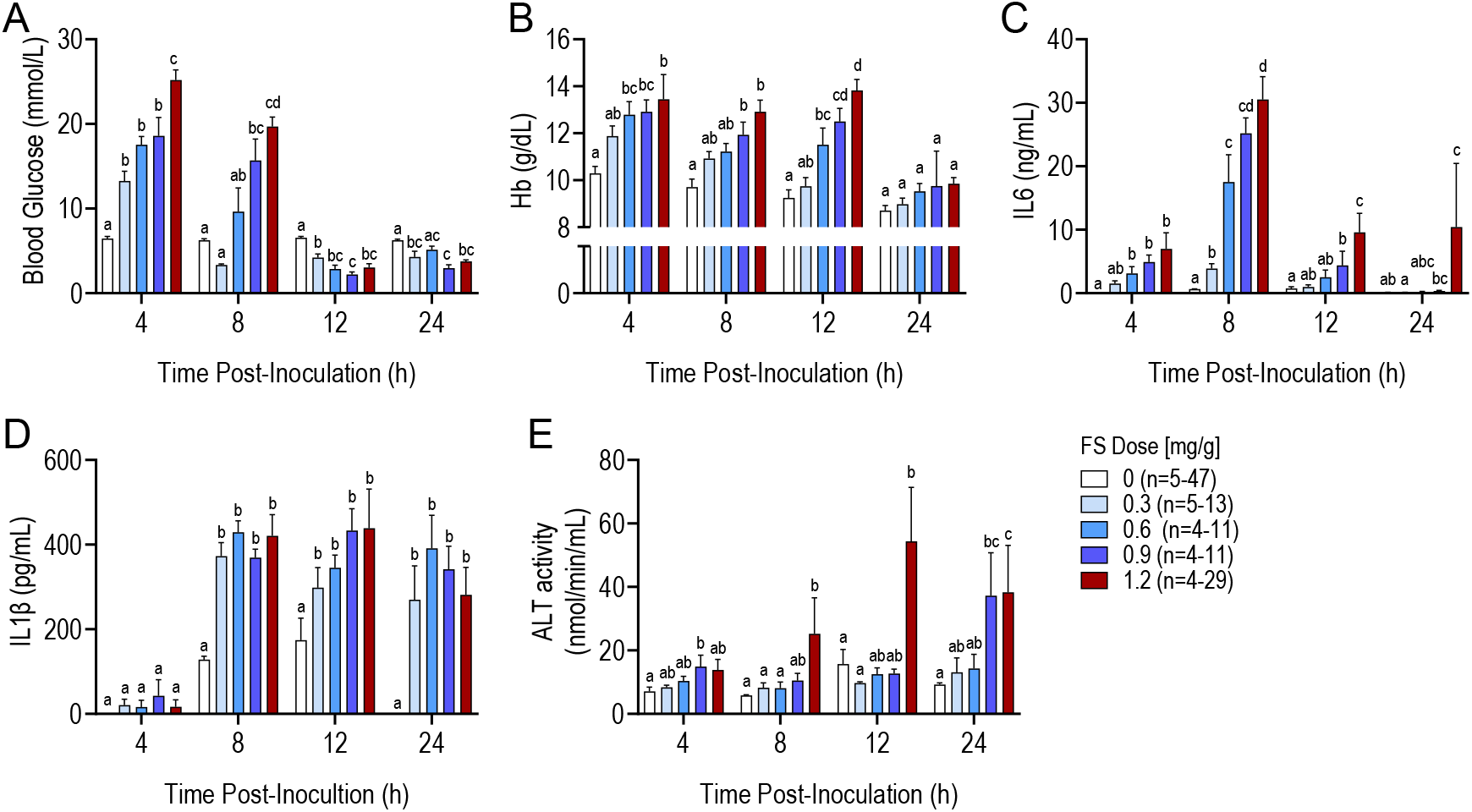
Dose- and time-dependent changes in (A) blood glucose, (B) hemoglobin (Hb), and plasma (C) IL-6, (D) IL-1β and (E) plasma alanine aminotransferase (ALT) activity in vehicle-and fecal slurry (FS)-challenged pups. Data is presented as mean±SEM. Dose effects on biochemical markers were assessed using linear mixed models, with sex as a fixed effect and dam as a random effect to account for litter clustering. Different letters above bars denote significance (P<0.05) by Tukey’s post hoc test.

Septic pups also exhibited acute and sustained cytokine release. IL-6 rose acutely and showed the steepest dose-response among all markers measured: at T8h, mean IL-6 reached 33– 35pg/mL in the 1.2 and 1.5mg/g FS dose groups compared to 0.7pg/mL in vehicle-treated pups (overall dose effect P<0.0001; **Fig. 4C**); 10 of 15 Tukey-adjusted pairwise comparisons were significant at this peak, with non-significant comparisons confined to the 0.6, 0.9mg/g BW) and the two highest doses (1.2 and 1.5mg/g BW) relative to one another. IL-6 declined toward baseline in lower dose groups by T24h but remained elevated in surviving 1.2mg/g FS dose pups (28 pg/mL). In contrast, IL-1β rose more slowly, such that no apparent change was evident at T4h (P=0.54) but was markedly increased by T8h and remained elevated through T24h (P<0.0001 at all later timepoints; **Fig. 4D**).

Plasma ALT increased dose-dependently in surviving pups, with the largest elevations at T12h in the 1.2mg/g FS dose group (mean 67 U/L vs 15 U/L in vehicle-treated pups; overall dose effect P=0.001; **Fig. 4E**), and remained elevated at T24 h (P<0.001), consistent with continued hepatocellular injury in animals that survived the acute phase. The High dose group lacked late-timepoint data because no 1.5 mg/g pups survived beyond T8h.

Notably, none of the biochemical markers showed significant sex-differences or sex × dose interactions (all P>0.10), in contrast to the female vulnerability observed in mortality and early nRSS.

### 3.5. Female vulnerability appears in clinical scores but not biochemistry or time of death

The female disadvantage observed in mortality (HR=0.53 in the Mid dose, HR=0.35 in the High dose; P=0.038 and 0.026, respectively) and in early nRSS scores (T4-T8h, all P<0.025) was not paralleled by sex differences at the biochemical level. Across all five biochemical markers measured, no significant sex differences were detected in surviving pups at any timepoint (all Mann-Whitney P>0.05; **Supplementary Table S3**), and survivor comparisons had sufficient statistical power to detect even modest effects (minimum detectable Cohen’s d ≈ 0.31–0.47 at 80% power). In non-survivors, biochemistry similarly showed no significant sex differences at the time of euthanasia (all P>0.10; **Supplementary Table S3**), albeit these comparisons were more modestly powered (minimum detectable d ≈ 0.81–2.19) due to smaller sample sizes. Time of death in non-survivors was identical between sexes (median 8.0h for both; P=0.83). Together, these findings raise the possibility that female pups in this model are not physiologically more vulnerable than males but rather exhibit more conspicuous behavioral manifestations of equivalent illness, leading to earlier crossings of nRSS-based humane endpoint thresholds and consequently higher apparent mortality.

**Suppl. Table S3.**
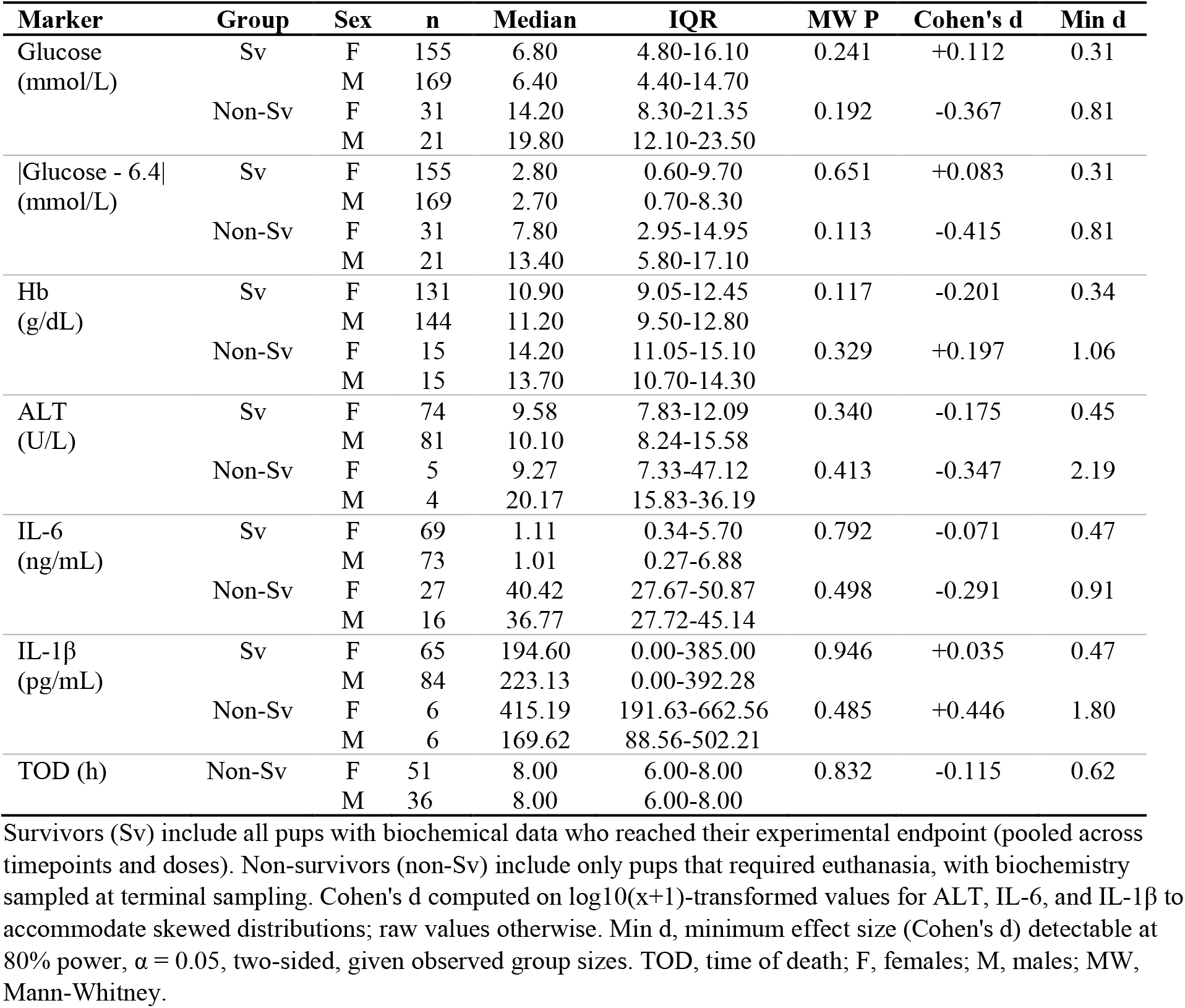
Sex-stratified biochemical comparisons in surviving and non-surviving pups.

### 3.6. nRSS validation against biochemical markers

To validate the newly developed nRSS, we examined its correlation with biochemical markers. For visualization, pups were grouped into nRSS bins (0–1, 2–3, 4–5, 6–7, 8–10; n < 3 were omitted; **Fig. 5A-E**), though correlation analyses of individual data points are shown in **Supplementary Fig. S2**. Among surviving pups, nRSS correlated significantly with all five biochemical markers measured (**Fig. 5A-E**; all P<0.0001). Glucose dysregulation magnitude rose monotonically across nRSS strata, from ∼2 mmol/L deviation in pups with nRSS 0–1 to ∼15 mmol/L in pups with nRSS 8–10 (Spearman ρ=0.70, n=270; **Fig. 5A**). Hb levels similarly tracked nRSS severity (ρ=0.48, n=224; **Fig. 5B**), as did IL-6 (ρ=0.54, n=131; **Fig. 5C**). Among the slow-resolving markers, IL-1β and ALT correlated with maximum nRSS (ρ=0.40, n=149 and 0.36, n=155 respectively; **Fig. 5D, 5E**). Across all five markers, bin-mean curves were monotonically increasing, indicating that nRSS faithfully captured a graded severity gradient consistent with underlying physiological disturbance.

**Figure 5.**
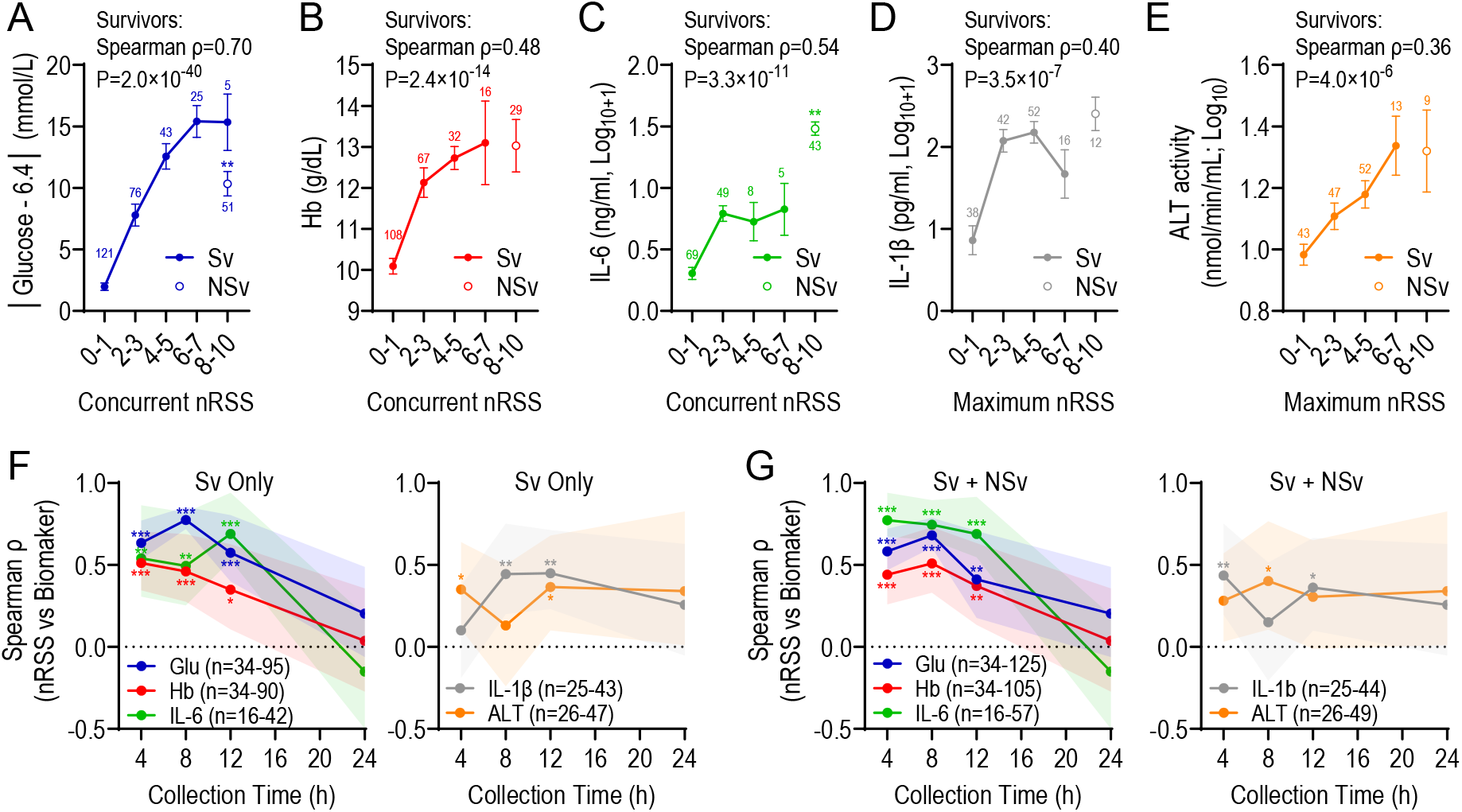
nRSS scores correlate with biochemical markers and capture the temporal evolution of disease severity. (A-E) Bin-means of biochemical markers across nRSS severity bins in surviving pups (closed circles), with non-survivors overlaid (open circles). Data in A depict glucose dysregulation, expressed as the absolute deviation from vehicle-treated median to capture the magnitude of disturbance regardless direction, due to the biphasic response observed. For panels A-C, nRSS is the score concurrent with biochemical sampling; for panels D, E, nRSS is the maximum score reached during the observation period, reflecting slower resolution kinetics of these markers. Data in A-E are mean±SEM; numbers above points denote n values. Cytokine data in C and D are plotted as log10(x+1) to accommodate zero values. Spearman ρ values and corresponding P values are computed on un-binned individual data from survivors (shown in Supplementary Fig. S2). Mann-Whitney U tests comparing non-survivors to nRSS 6-7 bin survivors: **P<0.01 (Glucose, IL-6); ns for all other markers. (F, G) Spearman correlations between nRSS and biochemical markers computer at each tissue collection time point, shown for survivors only (Sv; F) and for Sv and non-Sv (G). Shaded bands indicate 95% bootstrap CI (5,000 resamples). Asterisks denote timepoints where CI excluded zero: *95% CI, **99% CI, ***99.9% CI.

When non-survivors were overlaid at their maximum nRSS bin, they generally extended the survivor severity curve, as expected. For Hb, ALT, and IL-1β, non-survivor bin means lay close to or modestly above the rightmost survivor bin (**Fig. 5B, 5D, 5E**; Mann-Whitney P>0.16), confirming that nRSS captured biochemical disturbance continuously into the lethal range. For glucose, non-survivors showed lower magnitude of dysregulation than the most severely scored survivors (8.5 vs 17.9mmol/L; P=0.003; **Fig. 5A**), consistent with the biphasic glucose time course. That is, by the time pups reached lethal severity, the early hyperglycemic response had transitioned to terminal hypoglycemia, thus attenuating the absolute deviation. IL-6 was also a notable exception to the continuous severity gradient. Non-survivors had a median IL-6 of 39pg/mL versus 7pg/mL in the highest-severity survivors (nRSS 6-7), an ∼5-fold elevation above the survivor severity curve (**Fig. 5C**; Mann-Whitney P=0.004). Together, these data suggests that while nRSS accurately captures graded clinical severity across FS doses, certain inflammatory markers (i.e. IL-6) may have prognostic value beyond what is reflected in the clinical score alone.

To verify that the chosen nRSS score (concurrent vs maximum) reflected each marker’s resolution kinetics, we computed Spearman correlations between nRSS and each biochemical marker separately at each euthanasia timepoint (**Fig. 5F, 5G**). Glucose, Hb, and IL-6 showed strong correlations with concurrent nRSS at T4–12h (ρ=0.35–0.78) but lost correlation by T24h, when 77% of surviving pups had returned to nRSS=0, thereby leaving negligible variance in the score for correlation analysis. ALT and IL-1β, which remained at T24h, showed weaker but more stable correlations over time. These patterns confirm that nRSS reflects current physiological disturbance rather than cumulative illness exposure and validate the use of maximum nRSS for the slow-resolving markers in the primary analysis.

**Supplementary Figure S2.**
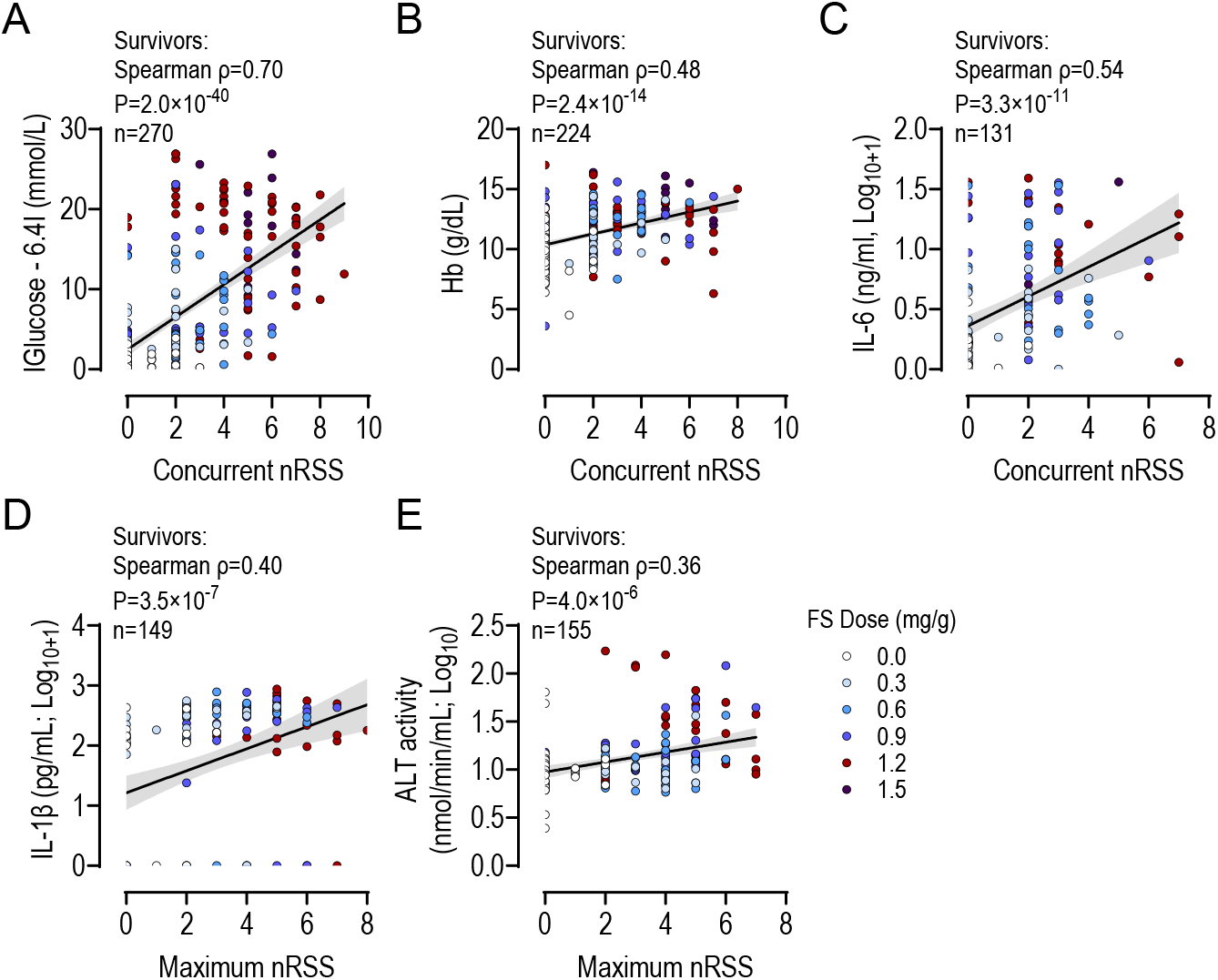
Individual level correlation between nRSS and biochemical markers. (A-E) Scatter plots showing individual surviving pups (one point per pup) colored by FS dose (0.3-1.5mg/g; vehicle controls also shown). Data in A depict glucose dysregulation, expressed as the absolute deviation from vehicle-treated median to capture the magnitude of disturbance regardless direction. For panels A-C, nRSS are plotted as concurrent scores (i.e. at the time of biochemical sampling); for panels D and E, nRSS are plotted as maximum score reached during the observation period. Lines and shaded bands represent linear regression with 95% CI for visual aid; reported statistics are Spearman rank correlations computed on individual data points. Cytokine data (C, D) are plotted on log10(x+1) values to accommodate zero values.

## 4. Discussion

Animal models are invaluable in biomedical research, providing a means to gain mechanistic insights into disease pathophysiology. However, the translational value of an animal model depends on the extent to which various characteristics, including intrinsic physiological differences (e.g. immune function, lifespan, developmental trajectories), environmental factors (e.g. housing, diet quality and quantity), and experimental design (e.g. age, primary and ancillary treatments, end-point selection), reflect the human condition being modeled. Among sepsis researchers, there is broad consensus that preclinical studies should adhere to rigorous design and reporting standards to improve the translation of preclinical findings. A key component of those standards is ensuring that animals are healthy at baseline and receive treatments consistent with clinical standards of care.

Here, we developed and characterized a neonatal rat FIP model in which pups were inoculated at PD3. As an altricial species, rats and mice are developmentally comparable at birth to preterm human infants (∼28-30wk of gestation),^32^ providing a relevant platform to study disease processes and the lasting effects of sepsis on immature organ systems. Increasing FS doses produced graded sepsis severity, reflected by increased inflammation, loss of barrier function (increased Hb), organ dysfunction (elevated ALT activity), and metabolic dysregulation (altered blood glucose). These physiological perturbations culminated in dose-dependent mortality, ranging from 0% to 100% by T24h. The mid dose (1.2mg/g) had an estimated mortality of 43%, which corresponds to upper thresholds of mortality reported in preterm and very low birth weight babies.^33,34^ Notably, this dose is higher than that required to induce comparable mortality in similarly treated adult mice,^35^ which at first glance appears at odds with studies showing neonates are more susceptible to bacterial and viral infections.^19,36,37^ This apparent paradox can be reconciled through a framework of disease tolerance versus disease resistance. Studies suggest that neonates mount an attenuated immune response,^19^ which limits their capacity to eradicate pathogens outright (disease resistance) but confers an enhanced ability to maintain host function despite high pathogen loads –a hallmark of disease tolerance. ^8,9,38^ In a disease tolerant host, a greater proportion of the administered inoculum would be expected to infiltrate the system, yet host function would be preserved at pathogen loads that would prove fatal in a disease resistant host. Importantly, neonatal pups in the present study received antibiotics as part of the modeled standard of care, which is not always applied in preclinical sepsis models.^19^ Without antibiotic therapy, neonates may be disproportionately affected by a higher pathogen burden and thus exhibit lower survival than their adult counterparts.^19^

A consistent feature of our model was the higher mortality observed in females compared with males at moderate and high doses, with females also showing higher early nRSS scores at T4h and T8h. By contrast, none of the biochemical markers measured differed between sexes, nor did any of the dose × sex interactions. Together, these findings suggest that the sex-dependent susceptibility to lethal sepsis in neonatal rats operates upstream of, or in parallel to the end-organ markers measured here, rather than through differences in the magnitude. The sex-differences in susceptibility to FS administration also explain the apparent protective effect of higher body weight on mortality: males were heavier than females in our cohort, and higher body weight tertile was disproportionately enriched for males, while the lower tertile was enriched for females. When sex and birth weight were considered jointly, the weight effect was abolished, indicating that body weight is unlikely to be an independent driver of survival in this model. The mechanisms underlying the female disadvantage at this developmental stage remain to be elucidated but may relate to sex-differences in innate immune programming, glucocorticoid signaling, or other developmentally regulated factors that influence the threshold for hemodynamic decompensation. Notably, the female disadvantage observed in our model contrasts with the male vulnerability typically reported in human preterm neonates,^39^ which may reflect species- or model-specific differences in the timing and direction of sex-dependent immune and endocrine maturation.

The second objective of this study was to develop a simple, sensitive, and rapidly administered nRSS to assess sepsis severity and monitor pup health over time. The ability to identify moribund pups promptly is essential for minimizing animal suffering and ensuring robust data collection through the consistent application of humane endpoints. Non-invasive scoring systems have been developed for neonatal septic mice,^30^ but differences in developmental milestones,^32^ body size, and litter characteristics between rats and mice necessitates species-specific criteria. Furthermore, the inclusion of clinical standard-of-care treatments (i.e. fluids, analgesics) could modify observable behaviours, and this had to be considered during nRSS development. At the outset, candidate criteria described in the published literature were considered.^30,40^ Several were found to be insufficiently reliable in pilot testing. Milk spot presence and skin temperature measurements were context-dependent (varying with time since last feeding and proximity to littermates, respectively) and therefore less informative. Breathing patterns and behavioural responses to tactile stimulation were difficult to assess consistently even with extensive training. Since speed and clarity of assessment were considered essential design criteria for the nRSS, these parameters were excluded from the final scoring system.

In contrast, skin color, righting ability, and regrouping behaviour worsened with illness severity and were therefore retained. A three-level color scale (pink, pale, grey) was straightforward to implement and reliably discriminated mild-to-moderate from severe illness in most cases, consistent with the observation in neonatal mice.^30^ Skin color assessment is feasible through the second postnatal week, after which fur growth limits visibility (though inspection of the ears, tail, face, and paws may extend its utility beyond this window). Righting ability was also a useful indicator of sepsis severity,^40^ though rat pups at PD3 differ from PD7 mice this respect: even healthy PD3 rat pups were periodically unsuccessful at righting, and righting times (10-50s) substantially exceeded the 4s time limit used in mouse studies.^40^ To accommodate this, rat pups were monitored for up to 60s and given an second attempt following a 2 min rest. This approach allowed assessment of the effort devoted to righting, rather than just categorical ranking of success or failure, which proved informative: vehicle-treated pups that failed to right invariably made vigorous and sustained attempts, in contrast to severely affected pups, whose effort tracked closely with overall health status. Finally, regrouping behaviour, assessed after pups were separated from the nest, was attenuated by FIP and provided a useful and complementary measure. The combination of all three parameters (color, righting, and mobility) achieved 8.5% overlap between survivors and non-survivors, compared with 15% for righting and mobility alone, and substantially higher overlap for any single parameter. The nRSS also demonstrate robust inter-observer reliability, supporting it practical utility in diverse research settings.

nRSS scores correlated significantly with all biomarkers of sepsis pathology examined, but the strength and timing of these correlations varied in ways that align with each marker’s resolution kinetics. Concurrent nRSS (i.e. the score recorded at the time of biochemical sampling) correlated robustly with markers that respond and resolve on a similar timescale to clinical signs, including glucose, Hb, and IL-6, the last of which is a commonly used biomarker for early diagnosis of neonatal sepsis.^41^ In contrast, IL-1β and ALT responded more slowly, but tended to remain elevated even after behavioural manifestations of illness had resolved, and these markers were therefore better captured by the maximum nRSS reached during the observation period. The delayed IL-1β response is consistent with its requirement for inflammasome priming and caspase-1 activation, processes that operate on a longer time scale than rapidly mobilized cytokines such as IL-6.^42^ Similarly, ALT activity was not appreciably elevated until T12h, consistent with the comparatively delayed onset of measurable hepatic injury. Importantly, the observation that nRSS correlated significantly with ALT as early as T4h, despite the absence of overt group differences in ALT at that time point, suggests the nRSS captures subtle gradations in sepsis severity that precede biochemically detectable organ injury – a valuable property for studies designed to evaluate early therapeutic interventions. Blood glucose exhibited a biphasic response: an initial dose-dependent rise followed by progressive decline into marked hypoglycemia at T12h-T24h, potentially reflecting early mobilization followed by depletion of glycogen stores (unpublished observations). Because of this directional change, glucose dysregulation was best captures as the absolute deviation from baseline, which preserved a strong correlation with nRSS across the observation period.

A further finding from the validation analysis was that, while nRSS captured a graded continuum of pathophysiology across most biochemical markers, IL-6 elevations in pups that eventually reached humane endpoints exceeded those of high-scoring survivors by approximately five-fold. A similar but less pronounced pattern was observed for IL-1β. This dissociation suggests that, although nRSS accurately reflects the global illness severity, certain inflammatory mediators, particularly IL-6, may carry additional prognostic information not fully reflected in the clinical score. This is consistent with clinical observations that very high IL-6 levels predict mortality in neonatal sepsis beyond what is captured by clinical scoring alone^41^ and reinforces the complementary value of biochemical and clinical assessment in both preclinical and translational contexts.

Monitoring schedules in preclinical sepsis studies should be calibrated to disease kinetics. In the present model, pups inoculated with higher FS doses showed a protracted illness trajectory that allowed close monitoring and timely identification of humane endpoints. However, a subgroup of pups, particularly those receiving moderate FS doses (0.6-0.9mg/g), exhibited an initial period of mild illness followed by rapid health deterioration. Based on these observations, we recommend nRSS assessments every 2h, beginning at T4h (when illness first becomes apparent) and continuing through T12h, after which mortality rates declined. For pups showing signs of severe illness at any assessment, increased monitoring frequency (e.g. every hour) is advisable.

Several limitations of the present model warrant discussion. First, the polymicrobial FIP model most closely approximates clinical scenario involving bowel perforation, such as those secondary to necrotizing enterocolitis, rather than the single-pathogen infections that predominate in neonatal sepsis (e.g. *staphylococcus epidermidis*, which accounts for >50% of late-onset sepsis cases in industrialized countries).^43^ The use of adult rat cecal contents as the FS source is cost-effective and yields consistent results, but whether the resulting microbial challenge is comparable to that produced by neonatal cecal contents is unclear. Nevertheless, adult-derived FS likely introduces a more complex polymicrobial inoculum, encompassing both gram-positive and gram-negative species, which may better reflect the clinical complexity of sepsis and may increase the likelihood that interventions demonstrating efficacy in this model will translate to a broader range of clinical presentations. Second, while PD3 is a well-justified developmental analog for preterm neonates at risk of sepsis, the generalizability of the nRSS to older postnatal ages has not been established. As pups mature, gain mobility, develop muscle strength, and open their eyes, behavioural responses to severe illness will change, and the scoring criteria propose here should be directly validated and, where necessary, modified for use at later postnatal timepoints.

An important caveat to the female vulnerability observation is that mortality in our model is, by design, triggered by nRSS-based humane endpoint criteria. To explore whether female pups were truly more physiologically affected, we compared biochemistry by sex in both survivors and non-survivors (Results, Supplementary Table S3). No significant sex differences were detected in any biochemical marker, nor did time of death differ between male and female non-survivors. These findings are consistent with two non-exclusive hypotheses. First, female pups may express behavioral manifestations of illness more conspicuously at equivalent underlying physiology, leading to earlier crossings of nRSS-based humane endpoint thresholds (behavioral expression hypothesis). Second, females may have a lower physiological threshold for translating equivalent biochemical disturbance into hemodynamic decompensation and mortality (lower decompensation threshold hypothesis), consistent with sex differences in cardiovascular reserve, autonomic regulation, and end-organ stress responses reported in other developmental contexts. Distinguishing these possibilities — and quantifying their relative contributions — would require severity measures uncoupled from the scoring system, such as direct cardiovascular monitoring or molecular markers of subclinical organ stress. More broadly, our findings highlight that sex effects in neonatal sepsis may operate downstream of routinely measured biochemical markers, with implications for both the interpretation of clinical scoring systems and the design of sex-stratified interventional studies.

In summary, we have developed and validated a clinically relevant neonatal rat FIP model that incorporates standards of care, along with a sensitive and reliable nRSS for longitudinal assessment of sepsis severity at PD3. The nRSS effectively predicts humane endpoints and correlates with established biomarkers of sepsis pathophysiology across the active phase of illness. Because early postnatal life represents a period of developmental plasticity, this model provides a platform not only for studying acute sepsis pathophysiology, but also for investigating the long-term health consequences of neonatal sepsis. The development of non-invasive tools such as the nRSS is therefore an important step toward enabling rigorous, reproducible, and welfare-conscious neonatal sepsis research.

## Acknowledgements

This work was supported by grants from the Canadian Institutes of Health Research (to SLB), from the Royal Alexandra Hospital Foundation (to KFM), and by the Stollery Children’s Hospital Foundation and the Alberta Women’s Health Foundation through the Women and Children’s Health Research Institute in the form of personnel awards to FJ (postdoctoral fellowship), SL (graduate studentship, summer studentship), KBT (summer studentship), and RMNN (graduate studentship). Additional funding for this project is provided by the Heart and Stroke Foundation of Canada and the Canada Brain Research Fund (CBRF), an innovative arrangement between the Government of Canada (through Health Canada) and Brain Canada Foundation (personnel award to JRR). SLB is a Canada Research Chair in maternal and perinatal physiology.

## Notes

### Competing Interest Statement

The authors have declared no competing interest.

